# Adjutant: an R-based tool to support topic discovery for systematic and literature reviews

**DOI:** 10.1101/290031

**Authors:** Anamaria Crisan, Tamara Munzner, Jennifer L. Gardy

## Abstract

**Summary:** Adjutant is an open-source, interactive, and R-based application to support mining PubMed for a systematic or a literature review. Given a PubMed-compatible search query, Adjutant downloads the relevant articles and allows the user to perform an unsupervised clustering analysis to identify data-driven topic clusters. Users can also sample documents using different strategies to obtain a more manageable dataset for further analysis. Adjutant makes explicit trade-offs between speed and accuracy, which are modifiable by the user, such that a complete analysis of several thousand documents can take a few minutes. All analytic datasets generated by Adjutant are saved, allowing users to easily conduct other downstream analyses that Adjutant does not explicitly support.

**Availability and Implementation:** Adjutant is implemented in R, using Shiny, and is available at https://github.com/amcrisan/Adjutant

## 1 Introduction

Literature reviews, whether systematic or not [7], are often the first step a researcher takes when entering on a new area of inquiry. Depending upon the subject, these reviews can involve hundreds or thousands of documents, which can easily become overwhelming, leading to significant culling of a document dataset to facilitate analysis [8]. This culling is antithetical to the purpose of the systematic review, yet few tools exist to help manage and explore these document datasets during the review process [10]. To address this issue, we developed Adjutant, an R-based tool supporting quick, visual assessments of topics within a document corpus. Like the military rank from which it draws its name, Adjutant’s purpose is to support an individual in their analysis process not supplement their domain knowledge.

## 2 Implementation Details

### Querying PubMed and assembling a document corpus

Given a PubMed-compatible search query, Adjutant uses the `RISmed package` [5] to obtain a summary of articles, including PubMed ID, Journal, Article Title, Authors, Abstract, Public Date, and MeSH terms. It then uses the `jsonlite package` [9] and the E-Utils eSummary API to query and extract additional metadata, including PubMed Central (PMC) ID, article DOI, PMC citation count, article type (i.e. Journal Article, Review, Meta-Analysis), and language. Both `RISmed` and `jsonlite` are used because of the differing outputs from the E-Utils eFetch and eSummary APIs.

### Data wrangling

Adjutant decomposes the PubMed document corpus into single-word entities extracted from article titles and abstracts, and converts it to a tidy format for further analysis using the `tidytext package` [11]. All words are stemmed using Porter’s algorithm [13], from the `SnowballC package` [1], after which common (stemmed) stop words are removed. Again using `tidytext package` resources, Adjutant next calculates the term frequency inverse document frequency (tf-idf) metric, and then filters terms that are too infrequent (fewer than 1% of all documents) or too frequent (more than 70%). Finally, Adjutant generates a document term matrix (DTM), with articles as rows, stemmed single words as columns, and tf-idf as the relevant analytic metric.

### Unsupervised Topic Clustering

The multidimensional DTM is decomposed into two dimensions using the Barnes-Hut t-SNE [12] implementation from the `Rtsne package` [6]. We use default t-SNE parameters, except when the document corpus contains more than 1000 articles and when the t-SNE the perplexity parameter is set to 50 [14]; however, Adjutant also allows users to modify the perplexity and theta t-SNE parameters after an initial analysis is complete. Next, Adjutant derives clusters using the hdbscan algorithm [2] from the `dbscan package` [4]. Adjutant will attempt to automatically calculate the optimal hdbscan minimum cluster points (minPts) parameter, with optimal being defined as the fewest number of clusters that best fits the t-SNE data. Adjutant identifies the optimal minPts parameters by leveraging goodness-of-fit measurements derived from linear models, specifically the adjusted *R*^2^ and the Bayesian Information Criteria (BIC); thus each minPts parameter value tested will have an associated *R*^2^ and BIC measure. Adjutant makes this calculation by fitting separate linear models to each of the two t-SNE dimensions, where for each linear model the t-SNE component co-ordinates are used as the dependent variable and the clusters are used as the independent variables. Each cluster is a vector of membership probabilities, from 0 (not in the cluster) to 1 (definitely a cluster member). The adjusted *R*^2^ between the two component models are multiplied, and the BICs are averaged. To choose the optimal minPts parameters, Adjutant identifies all minPts values with an adjusted *R*^2^ within 0.05 of the best performing minPts value, and among those different options selects the minPts value with the lowest BIC. The clusters resulting from the optimal minPts value are named using the two most commonly occurring terms within the cluster.

**Fig 1.**
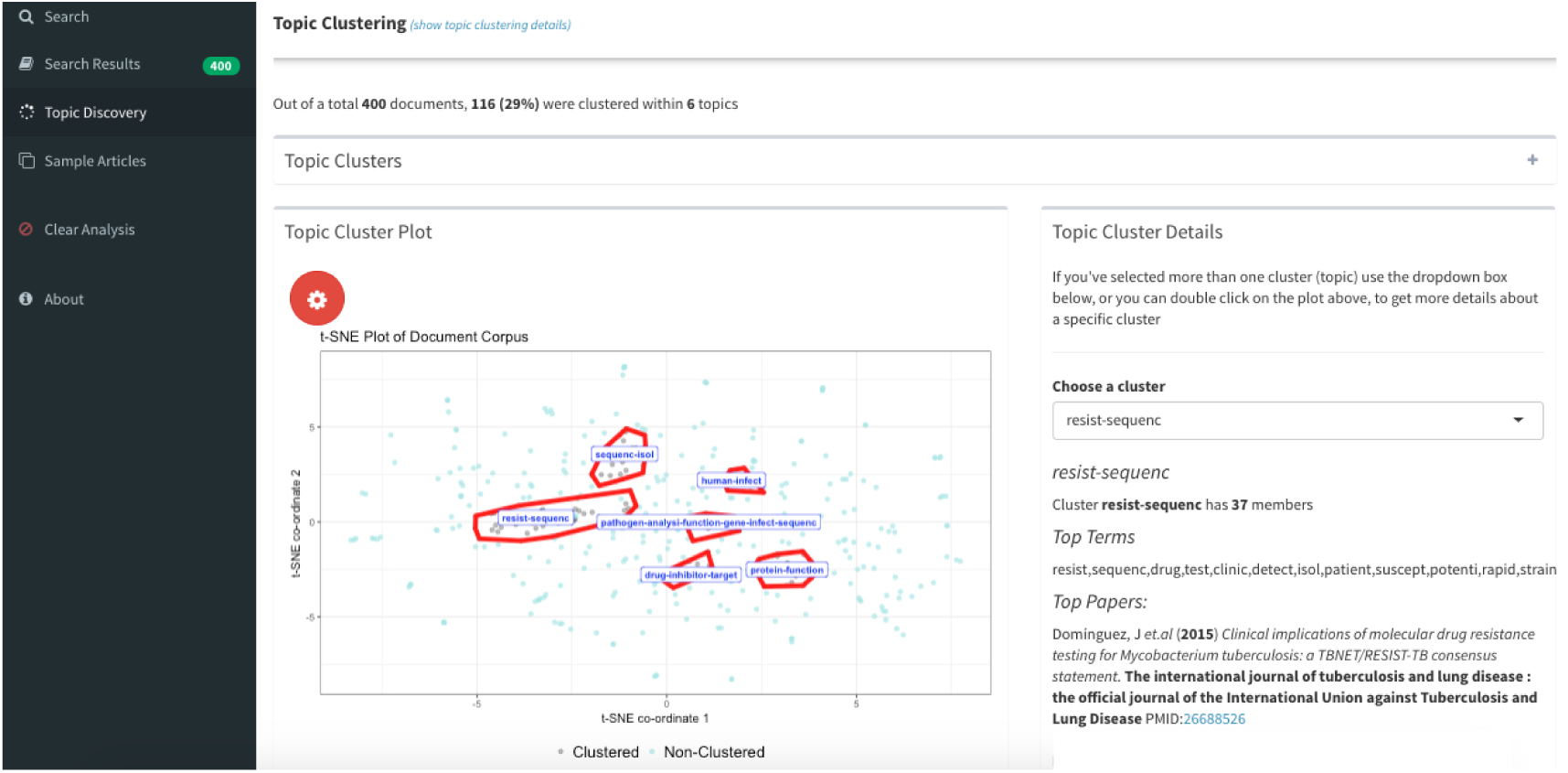
The Adjutant user interface after running the unsupervised topic clustering.

### Sampling

Adjutant uses the full document corpus for unsupervised topic clustering; however, a user may wish to produce and export a more manageable subset of the data for additional analyses. Adjutant allows users to filter their dataset and, if desired, to perform random sampling, including stratified sampling, with or without sampling weights.

### Using Adjutant

Adjutant is deployed as a Shiny application [3] within R, providing users with a graphical interface for performing search queries, inspecting the search results, initiating topic clustering, and sampling the dataset. Adjutant has a responsive interface that attempts to guide a user through the analysis steps. These steps can also be integrated into an R script, bypassing the Shiny application.

## 3 Conclusion

Adjutant is an R-based application that supports systematic and traditional literature reviews by enabling users to quickly visualize and explore the topic structure of a set of PubMed-derived documents. Importantly, by generating R-compatible outputs, Adjutant enables users to easily carry out downstream analyses beyond the scope of the tool.

